# Computational identification of surface markers for isolating distinct subpopulations from heterogeneous cancer cell populations

**DOI:** 10.1101/2024.05.28.596337

**Authors:** Andrea L. Gardner, Tyler A. Jost, Amy Brock

## Abstract

Intratumor heterogeneity reduces treatment efficacy and complicates our understanding of tumor progression. There is a pressing need to understand the functions of heterogeneous tumor cell subpopulations within a tumor, yet biological systems to study these processes *in vitro* are limited. With the advent of single-cell RNA sequencing (scRNA-seq), it has become clear that some cancer cell line models include distinct subpopulations. Heterogeneous cell lines offer a unique opportunity to study the dynamics and evolution of genetically similar cancer cell subpopulations in controlled experimental settings. Here, we present clusterCleaver, a computational package that uses metrics of statistical distance to identify candidate surface markers maximally unique to transcriptomic subpopulations in scRNA-seq which may be used for FACS isolation. clusterCleaver was experimentally validated using the MDA-MB-231 and MDA-MB-436 breast cancer cell lines. ESAM and BST2/tetherin were experimentally confirmed as surface markers which identify and separate major transcriptomic subpopulations within MDA-MB-231 and MDA-MB-436 cells, respectively. clusterCleaver is a computationally efficient and experimentally validated workflow for identification and enrichment of distinct subpopulations within cell lines which paves the way for studies on the coexistence of cancer cell subpopulations in well-defined *in vitro* systems.

## Introduction

Cellular heterogeneity within tumors contributes to resistance against chemotherapy and targeted therapy^1–3^, making it one of the most significant factors in disease relapse and mortality. Single-cell sequencing technologies continue to reveal heterogeneity within primary patient tumor cells and within *in vitro* model systems^4–7^. This intratumoral heterogeneity can be interpreted from an eco-evolutionary perspective in that subpopulations of cells adapt, interact, mutate, proliferate, and perish in response to their environment^8,9^. While single-cell sequencing technologies have been crucial to revealing the existence of novel cancer cell subpopulations, these assays are often endpoint. To understand and probe into mechanisms which drive tumor heterogeneity and tumor progression more fully, there is a pressing need for tools which allow identification, isolation, and perturbation of coexisting tumor cell subpopulations.

Multiple methods have been developed which aim to find minimal sets of marker genes which can be used to define groups, primarily aimed at tools which have limited gene panels^10–14^. Several other methods have aimed to find 1 or 2 surface marker genes which will maximally separate subpopulations^15–17^; only one of these methods, COMET, demonstrated experimental validation. Most of these approaches have relied on determining an optimal expression threshold built on the assumption that RNA expression will directly correlate with protein expression, but this is not always guaranteed to occur. A highly expressed marker gene is not guaranteed to be a highly expressed protein^18,19^, nor guaranteed to be localized to the cell surface in a particular experimental model^20^. Furthermore, optimal thresholds based on RNA expression do not have a direct conversion to flow cytometry. Finally, many of these methods are not easily scalable across large datasets, as they either require computationally expensive statistical methods or are not readily compatible with standard single-cell programs such as scanpy^21^ and Seurat^22^ which are optimized for the storage and analysis of complicated datasets.

To address these problems, we developed “clusterCleaver”, a computationally efficient and *scanpy* compatible workflow in which the Earth Mover’s Distance (EMD)^23,24^, a measure of statistical distance, is applied to rank individual surface marker genes based on how well they separate transcriptomic clusters of cells in scRNA-seq data. We preprocessed single-cell data to be better suited for predicting surface markers which could enable separation with FACS and developed it as a package compatible with the popular single-cell gene expression package s*canpy*. To test and validate the clusterCleaver workflow (Fig. 1), multiplexed single-cell RNA-sequencing (scRNA-seq) was first performed on 5 breast cancer cell lines to identify cell lines with highly distinct transcriptomic subpopulations. Then clusterCleaver was used on cell lines with distinct subpopulations to identify and rank candidate surface markers which could be used to physically separate transcriptomic clusters within each cell line. Top candidate markers were then experimentally screened with flow cytometry and subpopulations were FACS separated from the top hit for each. TagSeq, a bulk 3’ RNA-seq method^25,26^, and differential expression analysis was performed on isolated subpopulations, to assess similarity with transcriptomic identities of the targeted scRNA-seq clusters.

**Figure 1.**
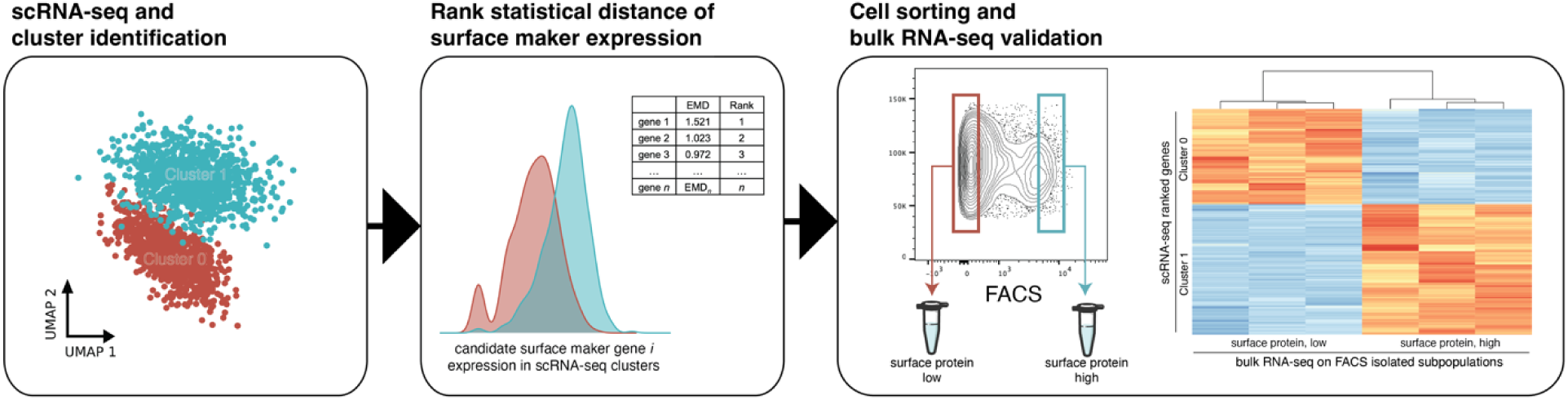
The clusterCleaver workflow. Taking clustered single-cell RNA-sequencing data as input, clusterCleaver computes EMD on predicted surface maker genes. Candidate surface marker genes are experimentally screened using flow cytometry, then surface staining antibodies which show subpopulation separation can be used to FACS isolate cell subpopulations. Transcriptomic identity of sorted cell subpopulations can be validated by performing bulk RNA-seq.

## Results

### Identification of subpopulations within cell lines through scRNA-seq

As previous studies have suggested the presence of multiple subpopulations within breast cancer cell lines^5,27–31^, multiplexed scRNA-seq was performed on 5 different unperturbed and early passage breast cancer cell lines (BT-474, MDA-MB-231, MDA-MB-436, MDA-MB-453, and Hs578T) to identify cell lines which contain distinct transcriptomic subpopulations (Fig. 2a). Leiden clustering was performed on each cell line (Supp. Fig. 1a, Fig. 2b-c), then Pearson correlation coefficient (PCC) was calculated between Leiden clusters for each (Supp. Fig. 1b). As clusters within MB231 (PCC=0.81) and MB436 (PCC=0.87) were the most dissimilar of the cell lines test (BT-474 (PCC=0.95), MDA-MB-453 (PCC=0.95), Hs578T (PCC=0.94)), MB231 and MB436 cells were chosen for testing and validation of the clusterCleaver workflow.

**Figure 2.**
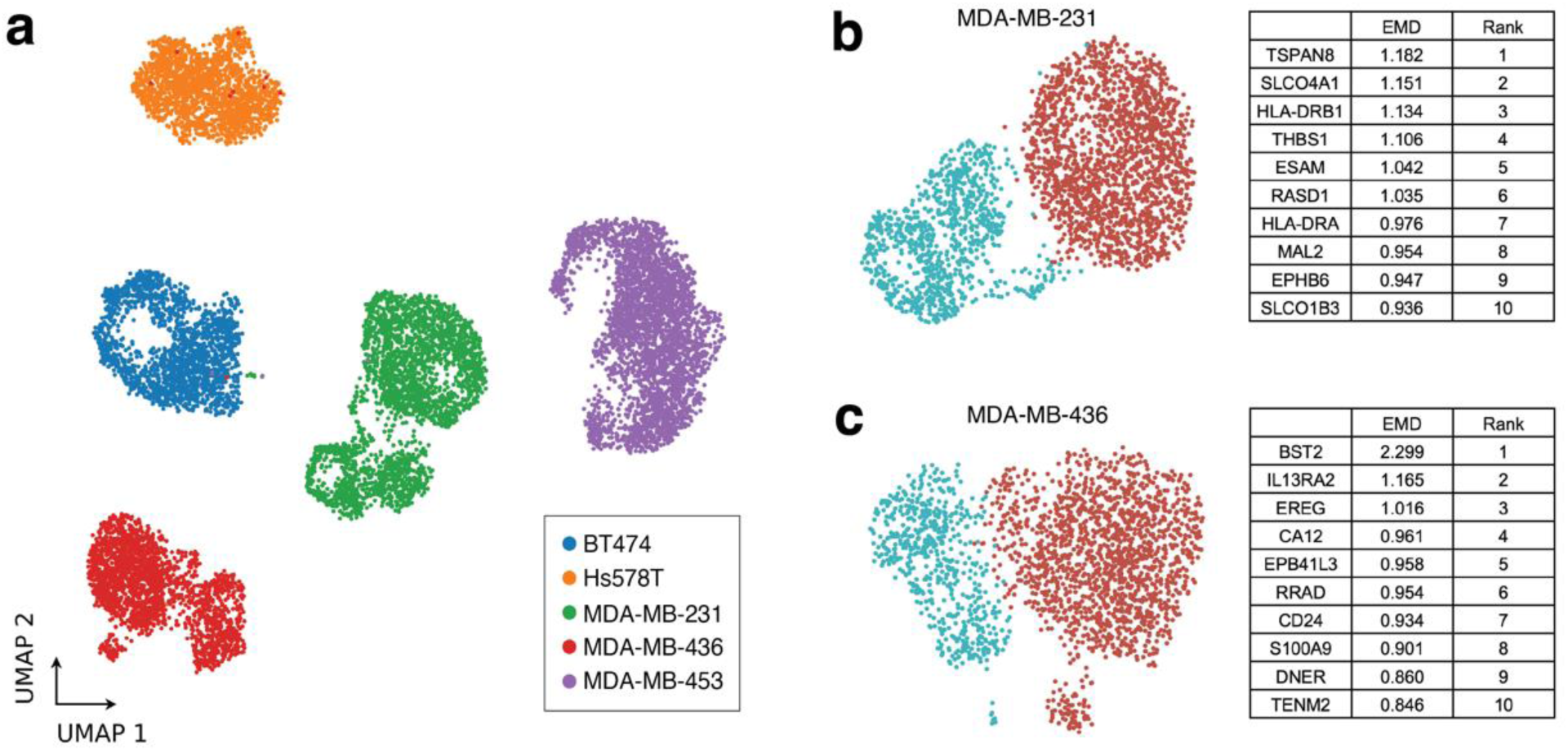
Identification of multiple transcriptomic subpopulations in commonly used breast cancer cell lines. (a) UMAP showing results of multiplexed scRNA-seq performed on BT-474, MDA-MB-231, MDA-MB-453, Hs578T, and MDA-MB-436 cells within 5 passages from ATCC. Leiden clustering and the top 10 EMD ranked candidate surface marker genes for separation of Leiden clusters for (b) MB231 and (c) MB436 cells.

### Application of the Earth Mover’s Distance

Next, we applied the Earth Mover’s Distance (EMD) to all genes within the scRNA-seq data ranked within the Cancer Surfaceome Atlas (TCSA)^32^ to identify candidate surface markers which could be used to physically separate unique transcriptomic clusters identified in scRNA-seq. EMD is a computationally efficient metric that compares two distributions. Intuitively, it can be thought of as the amount of work required to make two distributions equal^33^. Therefore, two distributions with low levels of overlap will have a relatively high EMD and highly overlapping distributions will have an EMD score close to 0. This property of EMD makes it highly suitable for quantitatively ranking marker genes in scRNA-seq with minimally overlapping and unique expression distribution patterns between transcriptomic clusters.

Given the goal of identifying genes that are likely to be expressed as surface proteins, EMD was applied to every candidate surface marker gene found both in the single-cell data and ranked within the TCSA database. TCSA provides a predicted surface score for each gene based on amalgamated data from nine different sources including experimental surface marker studies, computational protein conformation prediction, and prior database annotations^32^. TCSA surface marker scores are useful for filtering out genes unlikely to be surface expressed, but potential variation in mRNA translation and protein localization in different cell types highlights the need for experimental screening after running algorithms which depend on data built from other sources and cell types. The top candidate surface marker genes returned from clusterCleaver for MB231 and MB436 are summarized in Fig. 2b-c and a full list of EMD scores and rankings for each cell line can be found for each gene in Supplementary File 1 and Supplementary File 2.

### Screening of candidate surface markers

The top ranked genes for MB231 and MB436 from the clusterCleaver workflow with commercially available fluorochrome-conjugated monoclonal antibodies were screened (Supplemental Table 1). On MB231 cells, ESAM and TSPAN8 each identified distinct protein expression clusters by flow cytometry (Supp. Fig. 2) but when labeled in tandem, it was noted that TSPAN8 recognizes a subset of ESAM-high cells (Supp. Fig. 2a). HLA-ABC and ITGA2/CD49b provided positive surface staining, but without clear delineation between subpopulations (Supp. Fig. 2b-c). When stained in tandem with ESAM, HLA-ABC and ITGA2/CD49b showed expected cluster-specific protein expression patterns, with the ESAM-low subpopulation displaying higher average expression of HLA-ABC and lower average expression of ITGA2/CD49b as expected from scRNA-seq measurements (Supp. Fig. 2b-c). Tetherin (CD317), the protein product encoded by BST2, was found to be a surface marker for MB436 cells which identifies two distinct protein expression clusters by flow cytometry (Supp. Fig. 3). IL13RA2/CD213a2 and CA12 showed some positive surface staining (Supp. Fig. 3a-b), while antibodies against GYPC and EREG failed to stain MB436 cells (Supp. Fig. 3c-d). From results of this surface marker screen, we selected ESAM as the top flow cytometry candidate surface marker for separating clusters in MB231 cells, and BST2/tetherin for MB436.

### Isolation of subpopulations and validation of subpopulation transcriptomic identity

Returning to the goal of physically isolating subpopulations identified in scRNA-seq, we now asked whether the top protein surface markers identified using the Earth Mover’s Distance and screened with flow cytometry select for subpopulations with transcriptomic identities that match the respective scRNA-seq clusters. In the flow cytometry screen, ESAM (MB231, Fig. 3a-b) and BST2/tetherin (MB436, Fig. 4a-b) each revealed distinct subpopulations correlating to scRNA-seq cluster gene expression in their respective cell lines. FACS isolation was performed to enrich for ESAM-low/ESAM-high subpopulations of MB231 cells (Fig. 3c) and tetherin-low/tetherin-high of the MB436 cells (Fig. 4c).

**Figure 3.**
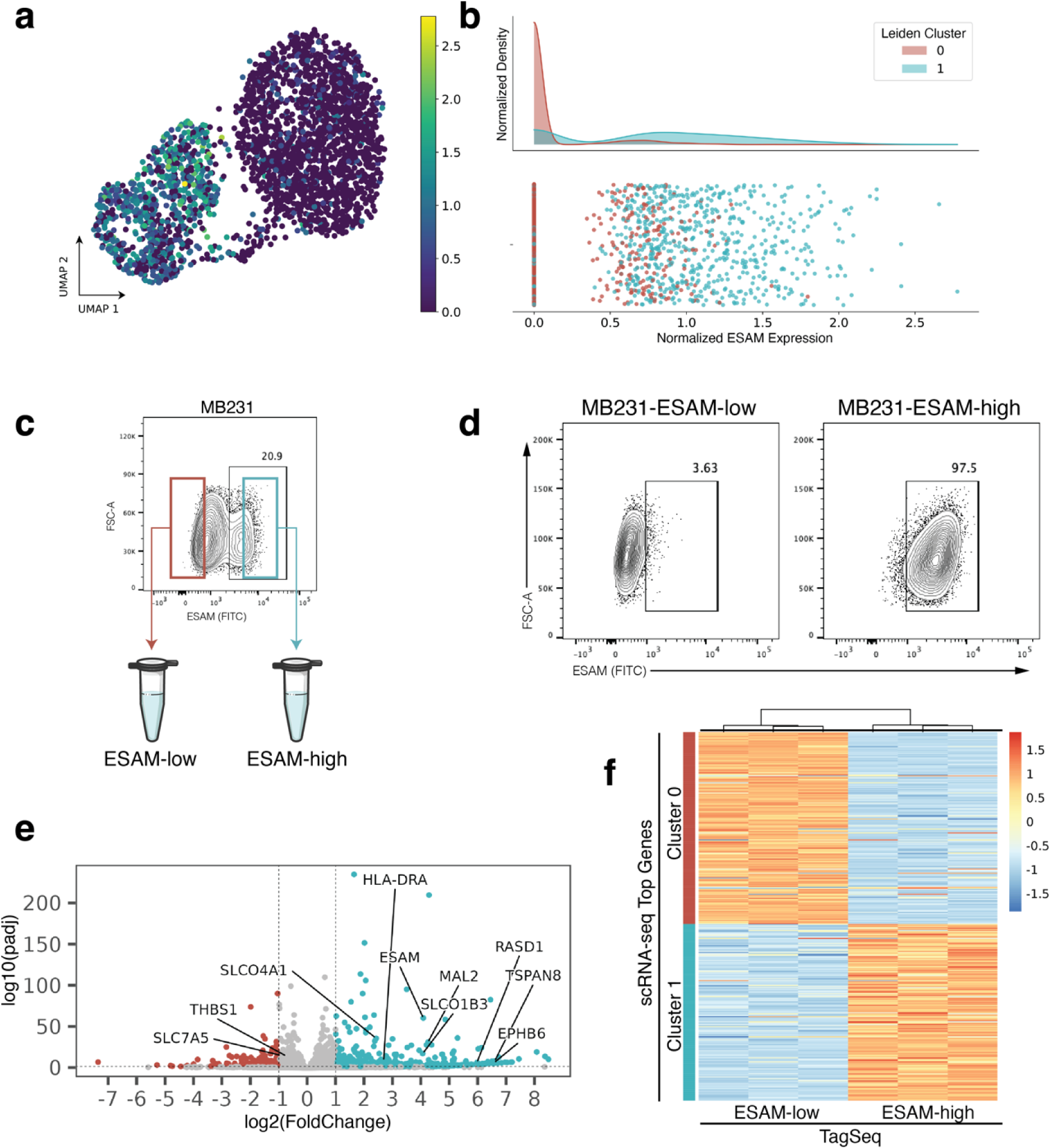
Experimental validation of ESAM as a surface marker for MB231 subpopulations. (a) scRNA-seq UMAP projection of MB231 cells colored by ESAM expression. (b) Histogram of ESAM expression showing enrichment of cluster 1 for ESAM. (c) ESAM staining on parental MB231 cells. Cells with the lowest staining were FACS enriched as “ESAM-low” and cells with the highest staining as “ESAM-high”. (d) At the time of RNA collection (10 days post-FACS and expansion), sorted ESAM subpopulations each maintained around 97% purity. (e) Volcano plot of bulk Tag-Seq performed on sorted subpopulations. Labels point to genes that were highly ranked in the scRNA-seq dataset using EMD. (f) Heatmap of bulk TaqSeq data for each FACS enriched subpopulation plotted against ranked cluster genes from scRNA-seq.

**Figure 4.**
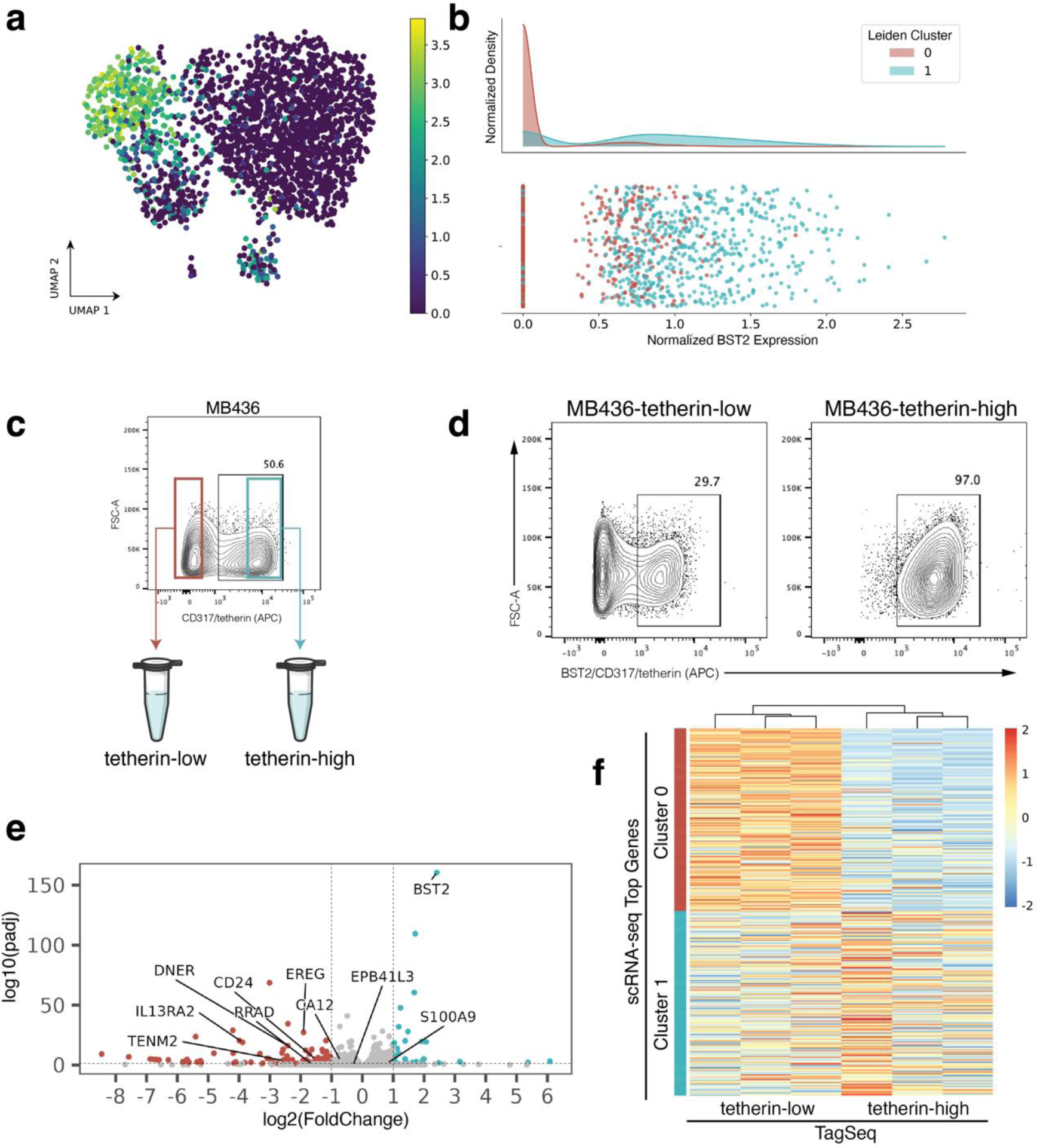
Experimental validation of BST2/tetherin as a surface marker for MB436 subpopulations. (a) scRNA-seq UMAP projection of MB436 cells colored by BST2 expression. (b) Histogram of BST2 expression showing enrichment of cluster 1 for BST2. (c) BST2 encodes the protein product tetherin/CD317. Tetherin staining reveals subpopulations within parental MB436 cells. Cells with the lowest staining were FACS enriched as “tetherin-low” and cells with the highest staining as “tetherin-high”. (d) At the time of RNA collection (13 days post-FACS and expansion), sorted the sorted tetherin-high subpopulation maintained 97% purity, however the tetherin-low subpopulation dropped to 70% purity. (e) Volcano plot of bulk TagSeq performed on sorted subpopulations. Labels point to genes that were highly ranked in the scRNA-seq EMD. (f) Heatmap of bulk TaqSeq data for each FACS enriched subpopulation plotted against ranked cluster genes from scRNA-seq.

Next, to check the transcriptomic identity of these isolated subpopulations, we expanded triplicate samples of each subpopulation after two rounds of FACS purification and then performed TagSeq, a bulk 3’ RNA-seq method^25,26^. At the time of RNA collection, parallel cells were immunostained and fixed to assess purity. We note that while the ESAM subpopulations of MB231 cells and the tetherin-high subpopulation of the MB436 cells maintained > 97% purity (Fig. 3d, Fig. 4d), the tetherin-low subpopulation of the MB436 cells were only 70% pure (Fig. 4d). The result has been repeated and is not due to sorting error. The inability of the MB436-tetherin-low subpopulation to maintain purity may indicate a unique biological property of these cells which suggests future studies.

To first assess if isolated subpopulations were significantly different from each other, differential expression analysis was performed on TagSeq data. Analysis with DESeq2 revealed 2250 differentially expressed genes (FDR < 0.05) in the ESAM-separated MB231 populations and 447 differentially expressed genes (FDR < 0.05) in the BST2/tetherin-separated MB436 populations (Fig. 3e, Fig. 4e).

Finally, transcriptomic identity of FACS isolated subpopulations was compared to transcriptomic identity of targeted scRNA-seq clusters. Top scRNA-seq cluster-specific genes were calculated by applying a t-test between scRNA-seq clusters. Comparing the TagSeq data of the FACS isolated subpopulations against the top cluster-specific genes from scRNA-seq, revealed that the transcriptome of the isolated subpopulations are well-matched to their expected scRNA-seq cluster identity (Fig. 3f, Fig. 4f). To quantify similarity between isolated subpopulations and scRNA-seq transcriptomic clusters, the top 50 differentially expressed genes for each subpopulation in TagSeq were then compared to the top 50 differentially expressed genes in scRNA-seq clusters. For MB231, 47/50 genes (94%) overlapped between the isolated ESAM-high subpopulation and its target scRNA-seq cluster (2/50 with the non-target cluster), and 46/50 genes (92%) overlapped between the ESAM-low subpopulation and its target cluster (3/50 with the non-target cluster). For MB436, 34/50 genes (68%) overlapped between the isolated BST2/tetherin-high subpopulation and its target scRNA-seq cluster (4/50 with the non-target cluster), and 38/50 genes (76%) overlapped between the BST2/tetherin-low subpopulation and its target cluster (5/50 with the non-target cluster). Despite differences in sequencing preparation and data generated from TagSeq versus scRNA-seq, we find a strong overlap of differentially expressed genes between isolated subpopulations and their targeted scRNA-seq cluster and weak overlap with the non-targeted cluster for each isolated subpopulation in both cell lines. These results suggest that clusterCleaver is a workflow which identifies surface marker genes which can be used to successfully enrich subpopulations from targeted scRNA-seq transcriptomic clusters.

## Discussion

We have shown that clusterCleaver is a computationally efficient workflow that takes in scRNA-seq and applies the Earth Mover’s Distance (EMD) to select surface markers which can enrich for targeted transcriptomic subpopulations from heterogeneous populations of cancer cells.

The use of EMD has been previously used as a method for separating flow cytometry data directly^39^, but also it possesses distinct advantages over other proposed methods for identifying markers for scRNA-seq data. EMD is computationally efficient^33^ and extensible to high-dimensional data^40^, making it ideal for searching across large scRNA-seq datasets. However, its primary advantage is that it does not rely on finding an optimal threshold or otherwise implementing a loss function which accounts for the sensitivity and specificity of a given threshold. EMD instead can be sensitive to outliers, meaning that a cluster with a subset of cells which have distinctly high expression of a given gene can potentially be highly ranked. This is advantageous for identifying markers in cases where a large bimodal distribution cannot be found. This does introduce scenarios in which a gene with a long-tailed distribution may be ranked higher than a gene with minimal overlap. To account for this, clusterCleaver includes several visualization modules to facilitate domain-specific interpretation when choosing surface markers to screen in flow cytometry.

All surface marker prediction methods are ultimately limited in that they are forced to rely on external databases to predict surface expression of genes and do not check for commercial antibody availability. Several computational methods have been developed which attempt to predict protein expression from scRNA-seq data^41–44^ which could be implemented to better search for markers which are more reliably expressed. clusterCleaver uses genes ranked in the Cancer Surfaceome Atlas (TCSA)^32^, but can be modified to rank custom surface gene lists or other sources as methods to predict surface expression of genes are improved.

clusterCleaver is not limited to use in cancer cell lines, however, we highlight this application for its implications for past and future *in vitro* studies. Numerous studies have been performed on cell lines without knowledge of their underlying population structure, yet subpopulations of cells may behave disparately under different culture conditions and perturbations skewing interpretation of bulk results. Subpopulations have been characterized in cell lines through different modalities^27–30,35,45,46^, but these studies may have isolated rare subsets of cells. In contrast, clusterCleaver delivers a workflow for discovering and isolating subpopulations starting from full knowledge of the transcriptomic diversity within cell lines by starting at scRNA-seq.

Currently, many studies which investigate eco-evolutionary dynamics in cancer *in vitro* rely on cocultures of cell lines from different genetic backgrounds^47,48^ or mixes of drug-naïve cells with lab evolved drug resistant strains^49^. While studies like these have been paramount in unraveling complex cancer dynamics, these cocultured subpopulations would not coexist together outside the lab. clusterCleaver is a workflow which can be used to identify flow cytometry markers to monitor populations changes in naturally coexisting cancer cell subpopulations within the same cell line and separate out subpopulations of cells which once coexisted together for further investigation. Using more physiological *in vitro* models of cancer cell heterogeneity may help reveal common mechanisms of coexistence and help guide improved therapeutic strategies for heterogeneous tumors.

## Methods

### Cell culture

All cell lines were used within 5 passages from thawed ATCC stocks at the start of this experiment. The passage numbers provided by ATCC via Certificate of Analysis are as follows: MDA-MB-231 (ATCC p31), MDA-MB-436 (ATCC p20), MDA-MB-453 (ATCC p349), Hs578T (ATCC p52), and BT-474 (ATCC p89). MDA-MB-231 and MDA-MB-453 were maintained in high glucose DMEM (Sigma, D5796) supplemented with 1X Penn-Strep (ThermoFisher, 15140122) and 10% FBS (Sigma, F0926). MDA-MB-436 and Hs578T were maintained in high glucose DMEM (Sigma, D5796) supplemented with 1X Penn-Strep (ThermoFisher, 15140122), 10% FBS (Sigma, F0926), and 10 µg/mL insulin (ThermoFisher, 12585014). BT-474 cells were maintained in Richter’s modified MEM without phenol red (ThermoFisher, A1048801) supplemented with 1X Penn-Strep (ThermoFisher, 15140122), 10% FBS (Sigma, F0926), and 20 µg/mL insulin (ThermoFisher, 12585014). FACS isolated subpopulations were maintained in their parental media. MDA-MB-453 were passaged with 0.25% Trypsin-EDTA (ThermoFisher, 25200056), all other cell lines were passaged with 0.05% Trypsin-EDTA (ThermoFisher, 25300062) using standard protocols. A cell scraper was used to fully release MDA-MB-436 cells after 1 minute of trypsinization.

### scRNA-seq sample and library preparation

Cells were prepared for multiplexed scRNA-seq using the 10X Genomics 3ʹ CellPlex Kit (10X Genomics, 1000261). Briefly, MDA-MB-231 (p4), MDA-MB-436 (p4), MDA-MB-453 (p5), Hs578T (p4), and BT-474 (p3) were gently detached, neutralized, strained through a 40 µm cell strainer, then counted. 1e6 cells from each population were added to a 2 mL tube and washed with room temperature PBS supplemented with 0.04% BSA (ThermoFisher, 15260037). Each cell line was resuspended in 100 µL of a unique cell multiplexing oligo and incubated for 5 minutes before 3 rounds of washing with cold PBS plus 1% BSA. After washing, cells were counted, and two pools were made containing equal ratios of 3 cell lines each: (Pool A) MDA-MB-231, MDA-MB-453, BT-474, and (Pool B) MDA-MB-436 and Hs578T. Cells were dropped off to the UT Austin Genomic Sequencing and Analysis and 15,000 cells per pool were loaded into a Chromium Next GEM Single Cell 3’ Chip following standard 10X Genomics protocols.

### scRNA-seq Analysis

Single-cell data was aligned to the GRCh38 (version refdata-gex-GRCh38-2020-A, 10X Genomics) and processed using cellranger’s (version 6.1.2) multi command. The data was then aggregated using cellranger’s aggr function. Cells were then loaded into scanpy (version 1.9.8). Quality control was done according to the scanpy’s single-cell best practices^34^. Automatic thresholding was done on ribosomal, mitochondrial, and hemoglobin genes using median absolute deviations (MAD) as defined by:

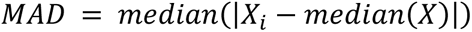

Where X is the expression of the given gene. Cells with a MAD greater than 5 were removed. Doublets were then removed using the package scDblFinder^35^. To normalize the data, we used the shifted logarithm technique as recommended by the single-cell best practices. Finally, we performed dimensionality reduction by finding highly variable genes, calculating the dataset’s principal components, computing a nearest neighbors graph, then calculating the UMAP representation. To remove cell cycle effects potentially biasing clustering, we regressed cell out cell cycle using gene lists from Tirosh et al^36^. All clustering and UMAP visualization was done using the Leiden algorithm as implemented by *scanpy* on regressed gene expression data. All candidate marker searches were performed on non-regressed log-normalized gene expression data. Similarity between clusters was computed between each cluster within each cell line on the full concatenated dataset by taking the top 50 principal component values of the gene expression and calculated the Pearson correlation coefficient between all identified clusters.

### Earth Mover’s Distance Pre-processing

From experience, ideal surface markers are bimodally distributed between clusters, with one cluster having gene expression around 0. However, one case that often occurs is that both clusters will have a peak of 0 gene expression while one of the clusters will have a second peak of higher gene expression. Therefore, this gene is potentially a good candidate as many cells from the high-expressing gene cluster can still be theoretically isolated. However, this gene will not rank highly with the EMD because the clusters both have significant overlap around 0. To circumvent this, we removed gene expression levels of 0 in clusters which had higher average gene expression. This allowed us to find more genes which were candidate markers.

### Flow cytometry and FACS

Cells gently detached and neutralized according to standard culture techniques. Cells were counted using a trypan blue exclusion automated cell counter. 0.2e6 cells were used for screening antibody labeling experiments. 1e6 cells were used for antibody validation and 2-5e6 cells were labeled for FACS cell sorting experiments. Cells were resuspended in cell staining buffer (PBS + 5 mM EDTA + 1% BSA + 1.6 mM NaOH + 0.01% sodium azide) and incubated in 1:100 diluted Zombie UV viability dye (Biolegend, 423107) or Zombie Violet viability dye (Biolegend, 423113) for 5 minutes on ice, then with the manufacturer recommended volume of antibody (antibody information in supplementary Table 1) for 20 minutes. Stained cells were washed 3 times with cell sorting buffer supplemented with 1:1000 diluted Zombie viability dye, then passed through a 40 µm cell strainer before flow analysis. For fixed cell preparation, cells were resuspended in 200 µL of 4% PFA in PBS for 15 minutes after the second wash, then washed 2 more times. Collection media for live cell sorting was prepared by supplementing complete media with 25 mM HEPES. Cells were sorted into 15 mL tubes containing 7 mL of collection media. Collected cells were spun down at 300 x g for 10 minutes, then plated in 50% conditioned media for 24 hours before transitioning to fresh, complete media. Conditioned media (CM) was prepared from complete media incubated on a 70% confluent plate of parental cells for 24 hours. CM was spun down at 500 x g for 10 minutes and supernatant was passed through a 0.22 µm filter. CM was diluted to 50% with fresh, complete media.

### RNA Collection for TagSeq

MDA-MB-231: MB231-ESAM-low and MB231-ESAM-high were FACS sorted 7 days before plating for RNA collection. 0.15e6 cells of MB231 p8, MB231 p13, MB231-ESAM-low, MB231-ESAM-high were plated across triplicate 6-well plates. Media was exchanged on plates after 24 hours. 72 hours after plating (10 days post-FACS), cells were 60-70% confluent and RNA was collected using an in-plate lysis strategy following the Qiagen RNAeasy Mini protocol. MDA-MB-436: MB436-tetherin-low and MB436-tetherin-high were FACS sorted 11 days before plating for RNA collection. 0.84e6 cells of MB436 p7, MB436 p13, MB436-tetherin-low, MB436-tetherin-high were plated across triplicate 6-well plates. Media was exchanged on plates after 24 hours. 48 hours after plating (13 days post-FACS), cells were 60-70% confluent and RNA was collected using an in-plate lysis strategy following the Qiagen RNAeasy Mini protocol. For all samples, RNA was eluted in 35 µL of nuclease-free water. RNA samples were quantified via Qubit, diluted, then submitted to the UT Austin Genomic Sequencing and Analysis Facility for 3’-Tag RNAseq (TagSeq) preparation.

### TagSeq analysis

Raw FASTQ files were processed using nf-core/rnaseq (v3.14.0)^37^ using default settings. Data was aligned to GrCh38. Bias uncorrected counts were rounded and used as recommended with DESeq2^38^ (v1.40.2) to generate differentially expressed genes and generate normalized expression values.

## Data availability

TagSeq data has been deposited in the Gene Expression Omnibus (GEO) under accession code GSE268250. Single cell data was deposited under accession code GSE268249. Flow cytometry fcs files are available upon request.

## Code availability

Notebooks used to process transcriptomic data and generate figures can be accessed at www.github.com/brocklab/clusterCleaver-analysis. The clusterCleaver package and instructions for installation and usage can be found at www.github.com/brocklab/clusterCleaver.

## Acknowledgments

We thank NIH F31CA268833 (to ALG), R01CA226258, R01CA255536, and U01CA253540 (to AB) which provided funding that supported this project and the training of personnel involved. ALG and TAJ thank Daylin Morgan for technical input on scRNA-seq analysis decisions. ALG thanks undergraduates Lan Zheng, Drew Saunders, Chisom Iloegbunam, and Andrea Ramirez who helped this project succeed by providing concurrent support to other projects while this project was taking place. 10X Genomics Single Cell 3’ Gene Expression and TagSeq preparation were performed by the Genomic Sequencing and Analysis Facility at UT Austin, Center for Biomedical Research Support (RRID: SCR_021713). Flow cytometry and FACS was performed at the Center for Biomedical Research Support Microscopy and Imaging Facility at UT Austin (RRID: SCR_021756).

## Author Contributions

**TAJ**: Conceptualization, Methodology, Software, Formal Analysis, Data Curation, Writing – Original Draft, Writing – Review & Editing, Visualization. **ALG**: Conceptualization, Methodology, Validation, Investigation, Data Curation, Writing – Original Draft, Writing – Review & Editing, Visualization. **AB**: Resources, Writing – Review & Editing, Supervision, Project administration, Funding acquisition.

