## Supplementary Information for "Computational identification of surface markers for isolating distinct subpopulations from heterogeneous cancer cell populations"

**Supplementary File 1. EMD ranked list of genes for MDA-MB-231 clusters.** Output ranked candidate surface marker genes from EMD for MDA-MB-231 scRNA-seq clusters. [*emdGenesRankScore231.csv*]

**Supplementary File 2. EMD ranked list of genes for MDA-MB-436 clusters.** Output ranked candidate surface marker genes from EMD for MDA-MB-436 scRNA-seq clusters. [*emdGenesRankScore436.csv*]

**Supplementary Table 1. Information on monoclonal antibodies screened on each cell line.**

| Cell Line | Gene | Antibody Target Protein | Cluster | Fluorochrome | Supplier | CatNo |
|---|---|---|---|---|---|---|
| mdamb231 | ESAM | ESAM | 1 | FITC | Miltenyi | 130-115-039 |
| mdamb231 | ESAM | ESAM | 1 | APC-Vio770 | Miltenyi | 130-115-038 |
| mdamb231 | TSPAN8 | TSPAN8 | 1 | PE | Miltenyi | 130-117-540 |
| mdamb231 | ITGA2 | CD49b | 1 | APC | Miltenyi | 130-100-328 |
| mdamb231 | SLC7A5 | LAT1 | 0 | FITC | R&D Systems | FAB10390G-100UG |
| mdamb231 | SLC7A5 | CD98 | 0 | PE-Vio770 | Miltenyi | 130-126-170 |
| mdamb231 | HLA-B | HLA-ABC | 0 | PE-Vio615 | Miltenyi | 130-130-114 |
| mdamb436 | BST2 | CD317/Tetherin | 1 | APC | Miltenyi | 130-101-660 |
| mdamb436 | CA12 | CA12 | 0 | FITC | Abcam | ab275576 |
| mdamb436 | GYPC | GYPC | 1 | Dylite405 | Novus | NBP2-33304V |
| mdamb436 | IL13RA2 | CD213a2 | 0 | PE | Biolegend | 360305 |

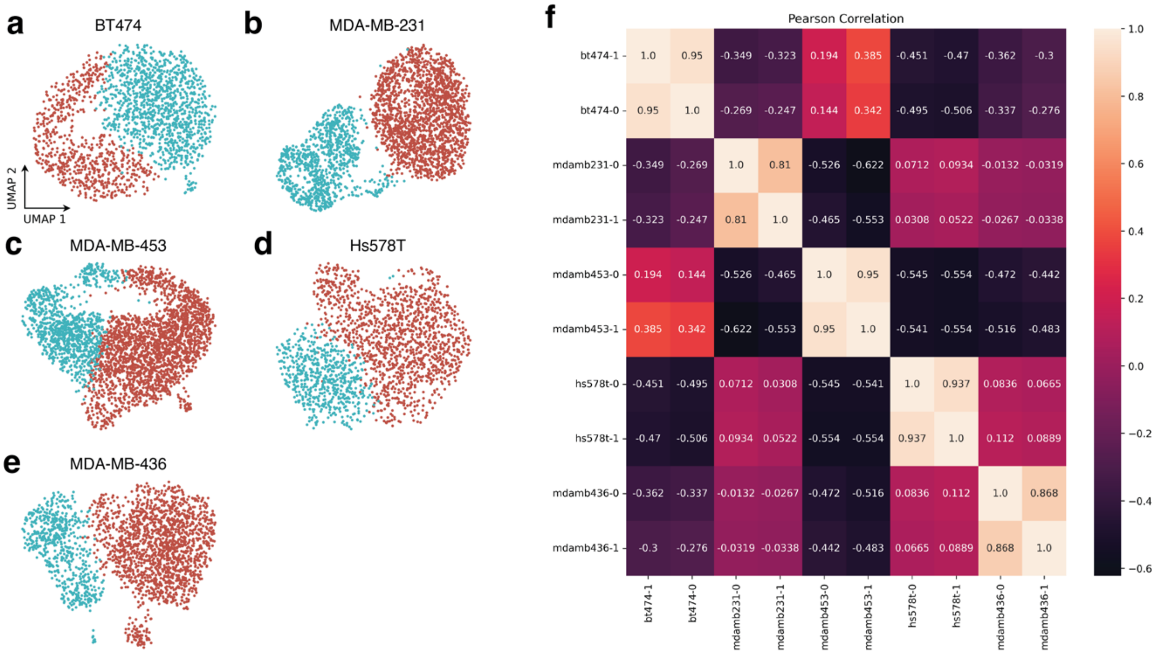


**Supplementary Figure 1. Clustering and cluster similarity within cell lines.** Clustered UMAPs for (a) BT-474, (b) MDA-MB-231, (c) MDA-MB-453, (d) Hs578T, and (e) MDA-MB-436**.** (f) Pearson correlation matrix showing similarity of clusters within and across cell lines.

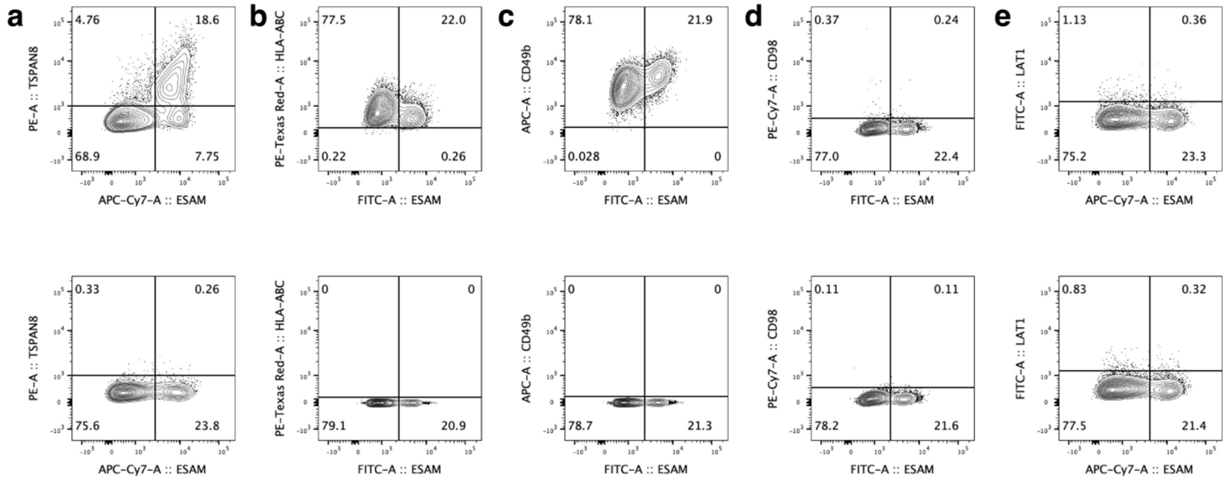


**Supplementary Figure 2. Screened antibodies on MDA-MB-231 cells shown as stained in tandem with surface marker, ESAM.** Top panel: double stained, Bottom panel: single ESAM-stained controls. (a) TSPAN8 stains a subset of ESAM-high cells, (b) HLA-ABC shows higher staining in a subset of ESAM-low cells, (c) ITGA2/CD49b shows higher staining in a subset of ESAM-high cells, (d) SLC7A5/CD98 did not surface stain, (e) SLC7A5/LAT1 did not surface stain.

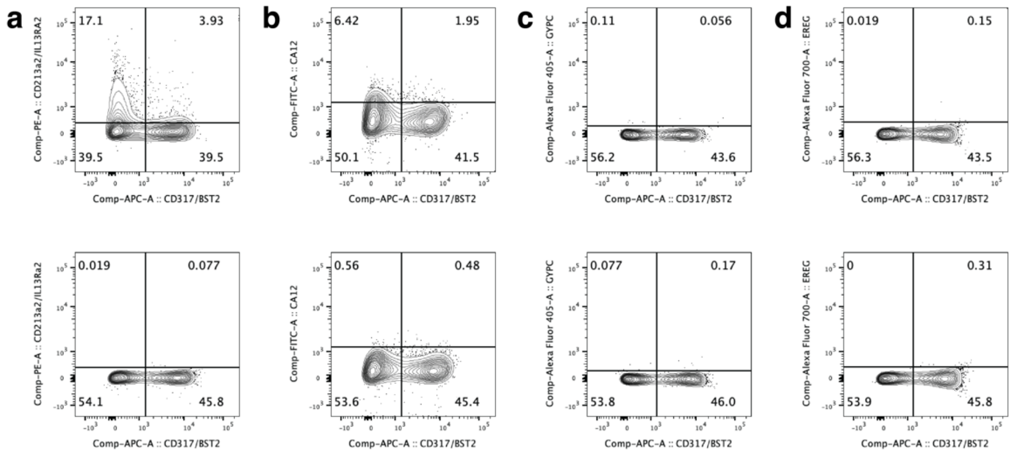


**Supplementary Figure 3. Screened antibodies on MDA-MB-436 cells shown as stained in tandem with surface marker, BST2/CD317/tetherin.** Top panel: double stained, Bottom panel: single BST2/CD317/tetherin-stained controls. (a) IL13Ra2/CD213a2 shows higher staining in a subset of BST2/tetherin-low, (b) CA12 shows higher staining in a small subset of BST2/tetherin-low cells, (c) GYPC did not surface stain, (d) EREG did not surface stain.
